# Evolved changes in DNA methylation in response to the sustained loss of parental care in the burying beetle

**DOI:** 10.1101/2021.03.25.436923

**Authors:** R Mashoodh, P Sarkies, J Westoby, RM Kilner

## Abstract

Levels of parental care critically influence the developmental environment with the capacity to impact the growth, survival, physiology, and behaviour of offspring. Plastic changes in DNA methylation have been hypothesised to modulate gene expression responses to parental environments. Moreover, these effects can be inherited and so may affect the process of adaptive evolution. In this study, using experimental evolution, we investigated how plastic changes in DNA methylation induced by the loss of parental care have evolved in a biparental insect (*Nicrophorus vespilloides*) using experimental evolution. We show that removal of care in a single generation is associated with changes in gene expression in stress-related pathways in 1st instar larvae. However, in larvae that have adapted to the loss of parental care after being deprived of care for 30 generations, gene expression is shifted from stress-related gene expression towards growth and brain development pathways. We found that changes in gene body methylation arose both as a direct response to the loss of parental care and stochastically as populations diverged. Overall, our results suggest that a complex interplay between transcription and DNA methylation shapes the molecular adaptation to environmental change.

## Introduction

There has been increasing interest in the idea that epigenetic mechanisms could participate in evolutionary change. Epigenetic mechanisms are the biochemical marks (*e*.*g*., DNA methylation, histone modifications) that ultimately determine how accessible DNA is to factors within the cell which, in turn, can regulate gene expression (1,2). These processes have been shown to be sensitive to the environment and represent a mechanism through which plasticity can be achieved during development. A hypothesised role for epigenetics in adaptation and evolution is driven primarily from recent work in plants and animals (*e*.*g*., *Arabidopsis, C. elegans*, mouse) demonstrating the capacity for transgenerational epigenetic inheritance *via* epigenetic modifications. These studies have shown that chronic challenges with stress, dietary changes or chemicals/drugs induce heritable changes in epigenetic marks in the germline that are transmitted to subsequent generations (3–6). The capacity for such inheritance and the degree to which it persists is highly dependent on the organism and the nature of the epigenetic change in question. Nevertheless, these observations raise the possibility that epigenetic mechanisms could buffer and/or modulate the pace of evolutionary change under changing environments. Indeed, theoretical models have supported this idea (4,7,8).

Although these ideas have generated much enthusiasm (and controversy), there is nevertheless limited empirical evidence to support the idea that epigenetic marks are induced by environments and persist in a way that drives long-term adaptive evolutionary change. There have been many observations of population-level differences in epigenetic marks between locally adapted wild populations in a range of diverse taxa (9–13) which are often correlated with local abiotic environmental conditions (*e*.*g*., salinity, temperature, latitude). The accumulation of methylation changes is suggestive of a potential role in adaptation to changing environments. However, there are a number of pathways through which epigenetic marks could participate in adaptative processes and contribute to the emergence of population-level changes. For example, epigenetic marks could be a by-product of transcriptional states or genetic sequence change (9,10,14). In this scenario epigenetic modifications are effectors and/or markers rather than drivers of the adaptive change. Taken together, epigenetic modifications could result from a complex interplay between plastic, genetic and stochastic processes acting in the short-and/or long-term (5). A critical first step towards understanding the sources of epigenetic variation is to characterise the extent to which epigenetic changes respond to an initial change in the environment, and the degree to which such changes might persist and be correlated with phenotypic changes (*i*.*e*., gene expression) as populations adapt to a changing environment.

Here, we used experimental evolution to examine how patterns of gene expression and DNA methylation are induced and evolve in populations of burying beetles as they adapt to the removal of post-hatching parental care (*i*.*e*., a change in the social environment). In natural populations of this locally abundant insect, burying beetle parents raise their young on a carrion nest (formed from a small dead animal, such as a mouse or songbird). There is continuous variation in the level of parental care provisioned (15). At one extreme, parents tend to their offspring throughout their development (including feeding with regurgitated pre-digested carrion) whereas at the other, parents abandon their offspring, leaving them to fend for themselves without any post-hatching care (16). Burying beetles are highly responsive to the levels of parental care they receive within a single generation (17). We have exploited this natural variation in care to establish two types of experimentally evolving populations in the laboratory (each replicated twice), which vary only in the family environment that larvae experience during development, and where the same family environment is created for successive generations within populations. In Full Care populations (FC_POP_), parents remain with their young throughout development; whereas in No Care populations (NC_POP_), parents are removed just before their offspring hatch (see Schrader et al., 2017; for more details on the experimental setup). Previous work has found that that NC_POP_ populations rapidly adapt to a life without parental care such that by generation 13 the proportion of successful broods matches that of the FC_POP_ populations (18,19). This adaptation to the loss of parental care is associated with a number of changes in morphology and behaviour. For example, NC offspring have evolved larger mandibles which likely helps with feeding on the carcass in the absence of parents (15), hatch more synchronously (20) and cooperate more with each other (21).

DNA-Me is a highly conserved epigenetic mark across plants and animals. This involves the covalent addition of methyl groups to CG dinucleotides (CpG) within DNA. DNA methylation (mCpG) is added by DNA methyltransferases (DNMTs 1-3; *e*.*g*., for maintenance during cell division and *de novo*) and, in mammals, can be actively removed by ten-eleven translocation enzymes (TETs) (22). Although parental care (and its loss) has been associated with plasticity in both DNA-Me and gene expression across a number of species (23–26), its role in insect DNA-Me and gene expression plasticity is still poorly understood. Burying beetles have a functioning DNA-Me system with levels of methylation that are comparable to other holometabolous insects and possess single copies of DNMT1-3 (27–29). These enzymes are highly conserved across multiple taxa and methylation occurs in actively transcribed genes suggesting important, yet still poorly understood, roles for DNA-Me in the species that carry this epigenetic modification (30,31).

In this study, we first characterized the changes in gene expression might underpin adaptation to the NC environment in developing larvae. To distinguish between evolved and environmentally-induced (plastic) changes, we used a common garden approach where offspring were exposed to the reciprocal parental environment (*i*.*e*., FC_POP_ had parents removed to create a NC environment or parents remained post-hatching in NC_POP_ to create a FC environment). We then asked if changes in DNA methylation are correlated with adaptive divergence (see Figure 1a for experimental design summary).

**Figure 1.**
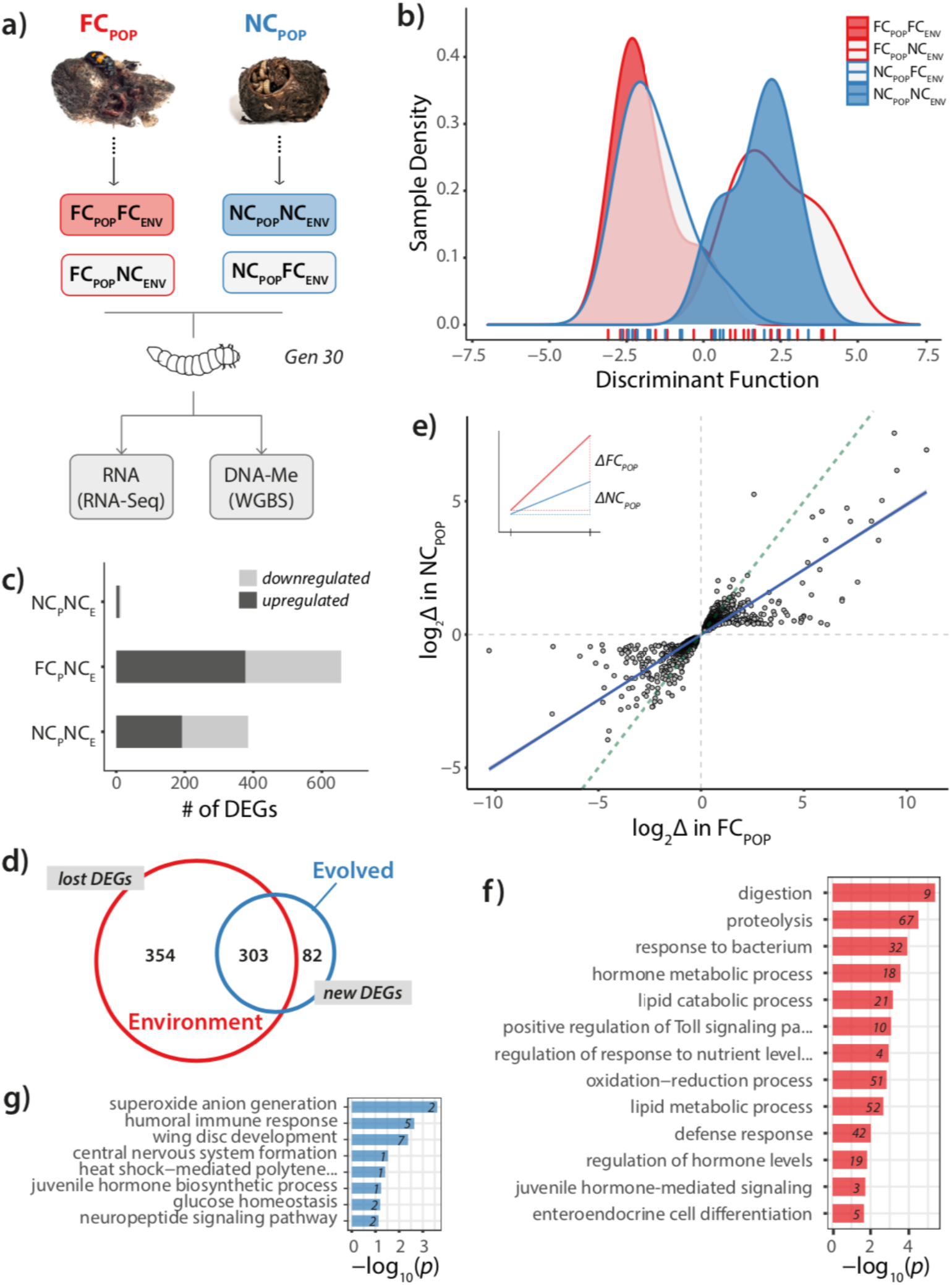
**a)** Summary of sampling from experimentally evolving populations and sequencing strategy. Larval heads from Full Care (FC_POP_) and No Care (NC_POP_) populations in native (FC_POP_FC_ENV_; NC_POP_NC_ENV_) and reciprocal environments (FC_POP_NC_ENV_; NC_POP_FC_ENV_) were sampled and used to measure gene expression (RNA-Seq) and DNA methylation (whole-genome bisulfite sequencing; WGBS). **b)** Density plots of discriminant function values for each family indicating that variation in gene expression is primarily due to the current social environment. **c)** Number of differentially expressed genes (DEGs) relative to the FC_POP_FC_ENV_ group. **d)** Venn diagram indicating degree of overlap between plastic and evolved DEGs. **e)** Scatterplot showing the correlation between log-fold changes (log_2_Δ) in the response to a NC_ENV_ within each population. **f)** Gene ontology (GO; biological processes) enrichment for plastic response to a NC_ENV_. **g)** GO terms for “new” DEGs following adaptation. Only GO terms with FDR-corrected *p*<0.05 are shown. Numbers indicate number of genes associated with that category.

## Results

### Differential gene expression in response to the sustained removal of care

To quantitatively characterise genome-wide variation in larval gene expression between our populations, we conducted a discriminant analysis of principal components (DAPC) using all expressed genes (n=12,772) within samples. The removal of care was significantly associated with a shift in gene expression (environment posterior mean = 4.22, 95% CI = [3.35, 5.15], *p*_MCMC_ < 0.001; Figure 1b) and this response was true whether individuals were derived from the FC or NC populations, as there was no effect of population (population posterior mean = 0.27, 95% CI = [-0.63, 1.20], *p*_MCMC_ = 0.576) nor an interaction between population of origin and the current environment (posterior mean = -7.91, 95% CI = [-2.12, 0.40], *p*_MCMC_ = 0.210). Further, each replicate responded to the care treatment in a similar way, since there was no effect of block on gene expression plasticity (posterior mean = 0.03, 95% CI = [-0.6, 0.72], *p*_MCMC_ = 0.940). Taken together, these results suggest that while there were large shifts in gene expression in response to the absence of parental care in the current environment, evolved changes in gene expression are likely to be more subtle and/or involve smaller subsets of genes.

To investigate these differences in more detail we performed differential expression analyses using FC_POP_FC_ENV_ as the reference population to estimate the number of differentially expressed genes (DEGs) across conditions. We found that 657 genes were differentially expressed when parental care was removed for the first time in the FC_POP_ (n=657; all log_2_ fold changes > 1; Figure 1c). By contrast, in the NC_POP_, which had experienced a NC_ENV_ for the previous 29 generations, 385 genes were differentially expressed which is a significant reduction in response (*X*^*2*^(3, n=12772) = 21558, *p*<2 × 10^−16^). Interestingly, only 20 genes were differentially expressed when NC_POP_ parents were allowed to care for their offspring for the first time in 30 generations, suggesting that gene expression was not constrained by their exposure to divergent selection pressures during these generations.

There was significant overlap in identity of the genes being differentially expressed in FC_POP_NC_ENV_ compared to the NC_POP_NC_ENV_ (log_2_(OR)=122.4265, *p*<2,2 × 10^−16^), with many more changes being lost (n=354) than being gained (n=82; Figure 1d). To test whether this reflected changes in the overall strength of response by individual genes to a NC environment within each population, we extracted overlapping DEGs (even those of small effect) and correlated the magnitude of response at each individual gene. This is the equivalent of comparing reaction norms on a genome-wide scale (See Figure 1e inset). We predicted that if the magnitude of change in response to a NC environment was similar across populations then we would expect fold-changes to be highly correlated resulting in a slope close to 1. We found that genes that were differentially expressed in response to a NC_ENV_ in both populations were positively correlated, with no evidence that genes switched direction of expression after evolving without parental care (slope = 0.489, *t*(3103)=91.875, *p*<2.2 × 10^−16^; Figure 1e). However, the slope of this correlation was significantly less than a slope of 1 (*F*(2,3103)=9176.4, *p*<2.2 × 10^−16^). Therefore, while there were similar changes elicited by a NC_ENV_ across both populations, the evolved NC populations expressed those genes at lower levels on average than the FC_POP_.

We extracted significant gene ontology (GO) terms from each contrast and collapsed across significant, but highly redundant terms, to look for overarching patterns. This analysis revealed that changes in gene expression primarily occurred in three broad categories of genes: 1) stress and its cellular response, 2) immune function and 3) growth and development (see Figure 1f for examples of GOs falling within these categories). Taken together, these results suggest that whilst the initial response to the removal of care is associated with an upregulated stress response and increased immune defense, individuals selected under a NC regime express fewer genes associated with these two categories. These changes are accompanied by enhanced expression of genes associated with physiological and neurobiological development (Figure 1g). Moreover, further examination of gene expression changes that emerge after 30 generations of exposure to a NC environment reveal an over-representation in the neurotransmission and neuropeptide categories (*e*.*g*., neuropeptide Y, orexin, and glutamate). For full list of GO categories see Supplementary Tables.

### CpG methylation in the burying beetle

We first sought to characterise the methylomes of first instar burying beetle larvae. We found average CpG methylation across the whole genome to be between 0.7-0.8% for all samples. Average levels of methylation were significantly higher in gene bodies (3.64 ± 8.60) compared to coding sequences (CDS; 2.72 ± 10.8), introns (0.952 ± 4.44), repeat/Tes (0.523 ± 1.96) and upstream regions (5’ UTR; 0.659 ± 3.22; all FDR-corrected Wilcoxon *p* values < 2.2×10^−16^; Figure 2a). Moreover, we found that gene body methylation was weakly but positively correlated with gene expression (*t*(1,11920)=36.34, *p*<2.2×10^−16^, *R*^2^ = 0.10). When we classified genes into expression state categories (silent, low, medium or highly expressed; Supp Figure 1a), we found that this correlation was mainly driven by expression states that were classified as medium or low. Medium and high levels of expression did not have significantly different methylation values from each other (Figure 2b, Supp Figure 1b).

**Figure 2.**
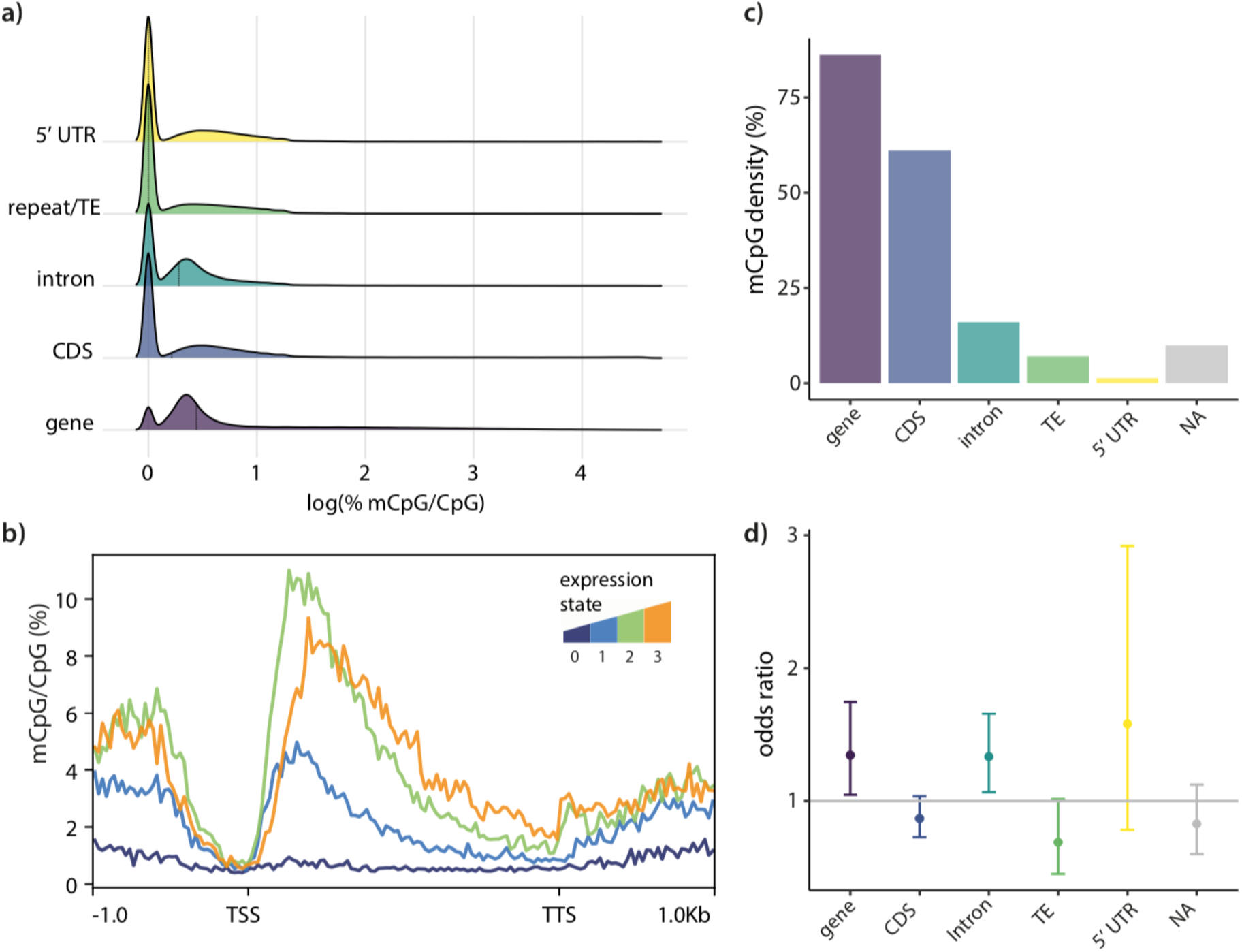
**a)** Ridge plots showing the distribution of percent methylated CpG levels (mCpG/CpG %) in different genomic features: 1000bp upstream of genes (5’ UTR), repeat or transposable elements (TE), introns, coding sequences (CDS) and across the entire gene body (gene). Median mCpG values are indicated by vertical dotted lines. **b)** Average mCpG levels across gene bodies (± 1kb) of genes categories by transcriptional activity: from inactive (0) to genes showing low (1), intermediate (2) and high (3) levels of expression. Plotted regions include 1kb upstream and downstream of transcription start sites (TSS) and transcription termination sites (TTS), respectively for each gene **c)** Bar plot indicating the distribution of methylated CpGs across gene features. **d)** Enrichment scores (odds-ratio; log_2_(OR)) for differentially methylated CpGs across different gene features. *\*Fishers’ FDR-corrected p<0*.*05*.

These findings are consistent with previous reports of methylation residing primarily in gene bodies for this species and the presence of higher methylation levels in gene bodies of genes in plants and animals (28–30).

Next, we determined whether the removal of care resulted in methylation changes in specific genomic features. We looked for enrichment of differentially methylated CpGs over the expected distribution of methylated CpGs across genomic features (Figure 2b). A CpG was classified as being methylated if the logit probability of it being methylated was greater than the bisulfite conversion error rate in at least 1 sample (see Methods). The density of methylated CpGs was higher in gene bodies (Figure 2c; *X*^*2*^(5, n=7400) = 14786, *p*<2.2×10^−16^) and we found differentially methylated CpGs (in both environmental and evolved contrasts) to be significantly enriched in genes and, specifically, introns (FDR-corrected *p*’s<0.05; Figure 1c) but not in any other annotated features tested. We, therefore, chose to focus on gene body methylation for subsequent differential methylation analyses.

### Differential gene body methylation in response to the sustained removal of care

We performed differential methylation analysis across gene bodies to compare (1) DNA-Me between populations (evolved changes; sustained exposure to a NC_ENV_ over 30 generations) with (2) DNA-Me induced by a single generation of exposure to a NC_ENV_ within the FC_POP_ (proxy for initial response) and (3) DNA-Me within the NC_POP_ when exposed to a FC_ENV_ (population-specific responses to NC_ENV_). As has been previously reported, not all genes in the burying beetle are methylated (28,29). Therefore, we clustered average weighted methylation values (calculated across the entire gene body), identified two distinct distributions of methylation states within genes (see Methods) and focussed our analyses on those that were classified as methylated (Supp Figure 2a, n = 4431; 39% of genes for which methylation was present for 85% of samples). These genes were also more highly expressed (W = 9161933, *p*<2.2×10^−16^, Supp Figure 2b) confirming that this procedure extracted functionally-relevant methylated genes. Consistent with previous findings, genes orthologous to housekeeping genes in *Drosophila melanogaster*, displayed higher methylation levels than those that were not housekeeping gene orthologs (*p<*2.2×10^−16^, Wilcoxon Test; Supp Figure 3).

We first compared the initial environmental response to a NC environment (within the FC_POP_) to the response 30 generations later (evolved differences). Removal of care in response to one generation of a NC_ENV_ resulted in 1486 differentially methylated genes, whereas we found 1530 differences when comparing the evolved response to the sustained loss of care (all genes FDR-corrected *p*<0.01). We also assessed the overall false discovery rate empirically by randomly shuffling samples between conditions 1000 times and extracting the number of differentially methylated genes (DMGs) after FDR correction at each shuffle. The number of DMGs detected in each comparison far exceeded the number that would be expected at random (*p*_FDR_ = 0 for both comparisons; Figure 3a). Moreover, the response to the removal of care was highly similar within each population (overlap of 741 DMGs; log_2_(OR)=1.8, *p*<2.2×10^−16^), though a high proportion of these changes remained specific to populations (Supp Figure 4). Taken together, this suggests that epigenetic variation can arise from environmental, population genetic background as well as an interaction between the two.

**Figure 3.**
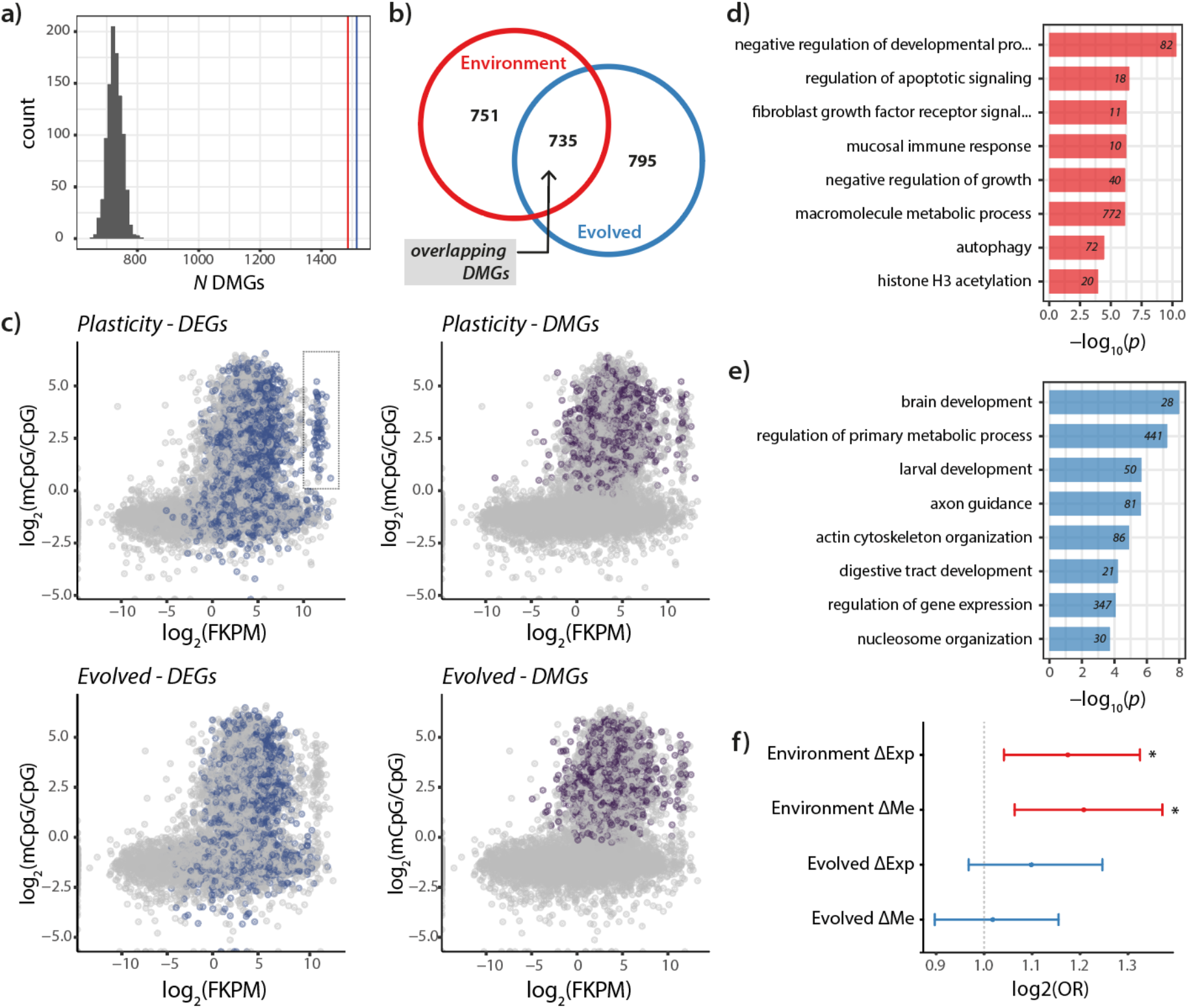
**a)** Histogram of number of differentially methylated genes (DMGs) passing FDR-corrected *p*<0.01 when samples are shuffled randomly. Vertical lines indicate number of DMGs found between plastic (red) and evolved contrasts (blue). **b)** Venn diagram indicating degree of overlap between plasticity and evolved DMGs. **c)** The relationship between gene expression (FKPM) and percent CpG methylation (mCpG/CpG) for all genes analysed for plasticity (top) and evolved (bottom) contrasts. Colours indicated differentially expressed genes (DEGs; blue) and DMGs (purple) for each contrast. **d)** Gene ontology (GO; biological processes) enrichment for environmental response to the loss of care DMGs and **(e)** Evolved DMGs. Only GO terms with FDR-corrected *p*<0.05 are shown. Numbers indicate number of genes associated with that category. **f)** Enrichment scores for associations between differential expression (ΔExp) and differential methylation (ΔMe) in housekeeping genes (* indicates corrected *p*<0.05).

### Differential gene body methylation and transcriptional change

Of the DMGs that were altered in response to a single generation of exposure to NC, 51% were significantly associated with differences in gene expression (all DEGs FDR-corrected *p*<0.01; log_2_(OR)=1.21, *p=*0.003). However, no significant association with gene expression was identified for responses measured after 30 generations of exposure to NC (log_2_(OR)=1.16, p=0.27). GO term enrichment analysis revealed that genes differentially methylated after exposure to one generation of NC were enriched in stress and growth reduction pathways (Figure 3d). Differentially methylated genes in the evolved response to a NC_ENV_ were enriched in biological processes related to growth and brain development (Figure 3e; see Supplementary Tables for a full list of GO terms). Furthermore, genes that were differentially methylated and differentially expressed in response to an environmental shift were significantly enriched (FDR-corrected *p*<0.05, Fisher’s Test; Figure 3f) for housekeeping genes compared to methylated genes. Interestingly, this enrichment was lost in genes that were differentially expressed in comparison between the two evolved populations (Figure 3f).

Given the high degree of overlap in DMGs induced by the environment compared to the evolved response to the NC_ENV_ 48%; Figure 3b), we hypothesised that the latter set of changes might reflect environmentally-induced changes that are no longer coupled with expression (i.e., methylation reflects ancestral rather than current transcriptional state). We, therefore, tested the overlapping DMGs against changes in gene expression induced by a single generation of exposure to NC. We found that overlapping DMGs are associated with the initial environmental response (log_2_(OR)=1.17, *p*<0.05) but not current changes in gene expression (log_2_(OR)=1.04, *p*=n.s).

To further investigate the idea that differential methylation can occur in evolved responses without associated differential gene expression, we highlighted differentially methylated genes on density plots describing the relationship between expression (FKPM) and average methylation for all genes. We identified a cluster of genes that was persistently differentially methylated but had lost differential expression after 30 generations of sustained exposure to NC (Figure 3c). Differential methylation in this cluster (outlined in Figure 3c) is associated with differential gene expression in populations exposed to NC for one generation (log_2_(OR)=1.91, *p =* 0.01) but not in the evolved response (log_2_(OR)=0, *p*=4.279e-05). Also differentially methylated at this cluster are genes from individuals in the NC_POP_ that were exposed back to the FC_ENV_ for a single generation suggesting they are environmentally labile (*i*.*e*., not inherited) (Supp Figure 5). We performed a GO analysis on this cluster and found these genes are almost exclusively enriched in ribosome biogenesis, protein translation and transport processes (see Figure 3f and Supplementary Tables).

We then asked if there was a possibility that environmental responses could become fixed evolved differences. In other words, DMGs that were initially responsive to the environment but no longer environmentally sensitive. We reasoned that such changes would not be reversed when NC_POP_ were provided with a FC_ENV._ There were 273 DMGs (18% of all evolved DMGs) that were differentially methylated in the evolved but not reverted by exposure of the NC_POP_ to a FC_ENV_. While these genes were enriched in defense, cellular and RNA metabolic processes these were not significantly associated with gene expression changes in larvae (log_2_(OR)=0.97, *p*=ns). This raises the possibility that some heritable methylation changes could become fixed during adaptive divergence, though the functional consequences of such changes remain unclear.

## Discussion

A role for epigenetic processes in mediating some of the molecular events underlying adaptive divergence is the subject of intense interest and debate (5,8,32,33). Debate persists because empirical studies testing the relationship between early environmentally-induced epigenetic responses and the subsequent adaptive evolution of such modifications to changing environments have been rare. How and when such changes are adaptive would depend on the persistence, heritability and function of epigenetic modifications during adaptation. Here, using experimental evolution in a wild-caught population of a subsocial insect, the burying beetle, we describe how gene expression and genome-wide patterns of DNA methylation diverge as beetles adapt to the experimentally imposed loss of parental care in the lab. We show that (1) DNA methylation and gene expression is highly sensitive to the loss of parental care, in both the initial and the evolved response (30 generations later), (2) The relationship between gene expression and DNA methylation at methylated genes shifts as populations diverge and (3) There is more variation in DNA methylation between populations (evolved change) and in population-specific responses to the environment than is reflected in gene expression profiles. Taken together, these data suggest that environmental, genetic and stochastic processes contribute to the accumulation of DNA methylation differences during adaptive divergence.

### Evolved changes in gene expression

Levels of parental care can shape the development of a number of physiological and behavioural traits that persist throughout the lifespan, including key fitness-related traits such as lifespan, fecundity and survival (17,26,34,35). Conversely, the removal of parental care has been shown to be associated with disruptions to behavioural development *via* changes in gene expression (24,26). Although much of the molecular level data comes from rodent model species, there is evidence for similar changes across primates, fishes and even a subsocial bee suggesting that there might be suites of conserved molecular mediators (23,25,36). Therefore, it is not surprising that the experience of a NC_ENV_ in burying beetles is the major source of variation in larval gene expression particularly in stress response pathways that disrupt growth and development. Adaptation to the loss of parental care in this species appears to be associated with the evolution of a stress-tolerant phenotype because we observe (1) reduced responsiveness of genes in the NC_POP_ to the NC_ENV_ (in terms of levels and number) and (2) the appearance of a small number of changes that may compensate for stress-related gene expression. This is consistent with the idea that adaptation to stressful environments and/or challenges involves fine tuning the balance between stress-and growth-related gene expression (López-Maury et al., 2008).

We further inspected the genes that were differentially expressed in the NC_POP_ but not FC_POP_ in response to the NC_ENV_ (*i*.*e*., new DEGs). One striking change was the high expression of a cytochrome P450 gene which appears to be a homolog to the Drosophila *Cyp6a20* gene, the deletion of which has been associated with higher levels of aggression and reduced sociality (37). Also, of particular interest was the differential expression of a number of odorant binding proteins. These proteins are odour sensing proteins that have been shown to be involved in a number of behavioural phenotypes. One such highly expressed gene was a homolog to Drosophila *Obp69a* which is involved in social responsiveness to social experience as well as starvation (38,39). Changes in the expression of both sets of genes was particularly interesting given that previous studies have shown that adaptation to the loss of parental care is associated with a greater level of coordination among larvae (18,20) and a shift from sibling conflict to cooperation (21).

### Sources of variation in DNA methylation

In rodent model systems, changes in gene expression in developing offspring in response to the loss of parental care have been shown to be, at least in part, mediated by changes in DNA methylation, most notably at promoter regions (24,36,40). Here we show that gene body methylation of developing larvae, though less widespread than vertebrate methylation, is similarly sensitive to the loss of parental care in a subsocial insect and that some of these changes are associated with differential gene expression. We further show patterns of DNA methylation diverge as populations adapt to the recurrent loss of parental care over multiple generations. Some of this change is simply induced afresh, each time the population is exposed to an NC environment. We know this because allowing parents to care for NC_POP_ offspring reversed some of these population-specific differences in DNA methylation. Nevertheless, some of the changes in DNA methylation that were induced originally by the environment did appear to become fixed over time in the NC_POP_. However, while a significant portion of DNA methylation resembles the initial response to the loss of care, many of the changes in DNA methylation that we observed were not explained solely by the removal of care. Some epigenetic changes are short-lived whereas others appear to persist suggesting that there might be functional differences in the timing and duration of such changes throughout the course of adaptation. These findings do not rule out the possibility of adaptive transgenerational inheritance of environmentally-induced methylation but they do suggest that population differences in epigenetic variation could emerge through multiple parallel processes.

Population differences in DNA-Me could be due, at least partially, to genetic variation that emerges through drift and/or selection. For example, in sticklebacks, some but not all population-level variation in DNA-Me is correlated with plasticity-induced by differences in water salinity (11). Some of these methylation changes have been linked to *cis* or *trans-*acting loci which show evidence of local adaptation (14). Similarly, population level changes in gene body methylation that appear in corals inhabiting different quality environments are associated with higher genetic differentiation at those loci (9). Another explanation is that population differences in DNA methylation could have arisen randomly (*i*.*e*., epimutations) and then been selected by the environment. This could be made more possible in insects by the reported high heritability of arthropod gene body methylation (12,41). Indeed, theoretical models have shown that stochastic but heritable epigenetic variation can be adaptive under changing environments (42). In plants, there is evidence for both stochastic, genetic and environmental sources of DNA methylation in explaining local adaptation (10,43,44). This further highlights the diverse mechanisms through which these changes in DNA methylation could emerge. Though we were able to distinguish between environmentally-induced and evolved changes in DNA methylation, this lays the groundwork for distinguishing between other (*e*.*g*., genetic, epimutational) sources of variation and its functional role.

Of particular note is that while a single exposure to the loss of care leads to an association between DNA methylation and gene expression, no such relationship is apparent after populations had diverged and the NC_POP_ had become adapted to a life without parental care. Indeed, over this timeframe, some methylation changes become uncoupled from differential gene expression, which suggests that adaptation to a NC_ENV_ might involve buffering against or overriding environmentally-induced changes in DNA methylation *via* other mechanisms. Our analysis of GO terms suggests that differentially methylated genes between the environmental and evolved conditions broadly reflect the same functional categories involved in divergent gene expression patterns. So while the initial response to the removal of care involves differential methylation in stress-related pathways, evolved differential methylation is enriched in processes associated with growth and brain development though this was not always associated with gene expression at the developmental point at which we measured it. DNA methylation in insects, including the burying beetle, appears to be stable across the lifespan (41,45). This raises the possibility that DNA methylation may be a product of ancestral transcriptional states. An alternative explanation is that these marks may prime transcriptional responses for activation at different developmental timepoints. What is clear is that while DNA methylation may not always reflect current transcriptional states, it accumulates as populations adapt and diverge and appears to be a marker for past environmental experience.

## Conclusions

There is accumulating evidence that populations accrue epigenetic variation that is correlated with local abiotic conditions, as they become divergently and locally adapted. These findings, coupled with work showing the transgenerational inheritance of environmentally-induced modifications (primarily in isogenic lab animals and plants) has provoked speculation that epigenetic modifications could play a key causal role in adaptation. An implicit, yet mostly untested, hypothesis here is that epigenetic modifications are inherited, and sustain adaptive divergence through a form of “plasticity-led” evolution. Here, we have taken an experimental approach to test this hypothesis. By manipulating environmental conditions (*i*.*e*., the loss of parental care) we have found that a significant amount of variation in DNA methylation can be induced and this is associated with transcriptional activity. However, after several generations of exposure to the same environmental cues, we find that population-level variation and transcription is not solely explained by the environmental differences. These indicate that additional transcriptional and stochastic processes contribute to the accumulation of differential DNA methylation as populations adapt and diverge. In short, although it is still possible that epigenetic variation contributes to adaptation, existing formulations of how this might work in practice are too simplistic to account for the experimental results we obtained.

## Methods

### Breeding design & Experimental Evolution

We analysed experimental populations of *Nicrophorus vespilloides* that had been evolving under different regimes of parental care, and which were founded from a genetically diverse founding population generated by interbreeding beetles from multiple wild populations across Cambridgeshire. These populations are described in detail in Schrader et al., 2017. and comprise a total of 4 populations, two blocks (Block 1 and Block 2; separated by 1 week) containing two populations evolving with (FC_POP_) or without parental care (NCpop). On the 29^th^ generation, seventeen days after their emergence as adults, when individuals were sexually mature, we paired 15 males and females within each population (N=30 pairs in total). Each pair was placed in a separate breeding box. Each pair was placed in a separate breeding box with moist soil and a thawed carcass (10-12g). We then placed each breeding box in a cupboard, and allowed parents to prepare the carcass and for the female to lay the clutch of eggs. After 53h, populations were split such that both parents were either removed (in keeping with the procedure experienced by the NC_POP_) or left in the breeding box. This produced offspring from both populations in their evolved condition (FC_POP_FC_ENV_ & NC_POP_NC_ENV_) as well as their reciprocal environments (FC_POP_NC_ENV_, NC_POP_FC_ENV_). At 80h post-pairing (approx. 10h post-hatching), first-instar larvae were collected from surviving families. Only families that with brood sizes between 20-30 larvae were used for subsequent RNA-sequencing analyses.

### Larval tissue dissection, RNA extraction & Sequencing

For each family, RNA from four heads of first instar larvae were pooled and extracted using Trizol (Invitrogen). Total RNA quality was checked using the BioAnalyzer System (Agilent), and yield was quantified using a Qubit RNA Assay Kit (Thermo Fisher). PolyA-selected RNA libraries were constructed and sequenced (150bp paired-end) at a depth of 30x by Novogene (Hong Kong). The resulting sample group sizes for gene expression analysis were: 5-6 libraries for each group within each block (total of 46 libraries).

### RNA Sequencing & Read Mapping

Reads were trimmed using TrimGalore (0.5.0; https://github.com/FelixKrueger/TrimGalore) to remove adaptor sequences, perform quality trimming and discard low-quality reads. Reads were mapped and quantified using a custom pipeline using HiSat2 (2.1.0) and Stringtie (2.0.3; Pertea et al., 2016). Transcript abundance estimation was based on counting reads aligned to the *N. vespilloides* reference transcriptome (NCBI Refseq Assembly: GCF_001412225.1; Cunningham et al., 2015). Lowly expressed genes (those with less than 15 counts in more than 90% of samples) were filtered from raw counts table leaving a total of 12,772 expressed genes in the dataset to be analysed.

### DAPC using adegnet and MCMCglmm

To compare overall patterns of gene expression plasticity, we performed a discriminant analysis of principal components (DAPC), which linearizes principal components within a dataset into a single discriminant function, enabling the comparison of overall levels of variation between groups. Gene counts were normalized and log-transformed using a regularized log transform in DESeq2 (version 1.26.0; Love et al., 2014) and a discriminant function was built by defining each populations response to their reciprocal environments using adegenet (version 2.1.3; Jombart, 2008). To compare differences in gene expression plasticity between populations we used the MCMCglmm package (version 2.29; Hadfield, 2010) to model DAPC score as a function of population background and current environment (and an interaction between the two) using a default prior that assumes a normal posterior distribution with large variances for the fixed effects and a weakly informative (flat) prior.

### Differential expression analysis

Differential expression was analysed using DESeq2 using custom scripts in R. Log2-fold change estimates for each differentially expressed gene (DEG) were shrunk to generate more conservative estimates of effect size using the ashr shrinkage estimator (50). Moreover, to focus on genes that are differentially expressed at higher thresholds we considered a gene to be significant differentially expressed only if it had increased by a 2-fold change (*i*.*e*., Log_2_FoldChange >= 1). We included the separate blocks as a covariate in all the analyses to increase power but also account for minor fluctuations in gene expression across the blocks of replicate populations.

### Functional Annotation

Functional enrichment analyses were conducted using the topGO R package (version 2.38.1; Alexa & Rahnenfuhrer, 2016) to identify over-representation of particular functional groups within the DEGs in response to the removal of care as well as the evolved response to the removal of care, based on GO classifications using Fisher’s exact test. GO terms were annotated to the *N. vespilloides* genome using the BLAST2GO (version 5.1.1) workflow to assign homologs to the *Drosophila* non-redundant protein databases (51). To increase functional predictions of *N. vespilloides* genes these annotations were supplemented with GO term assignments based on ortholog searches within Arthropods using eggNOG-mapper (52) and OrthoDB (v10.1; Kriventseva et al., 2019).

### Bisulfite Sequencing & Read Mapping

For each family, DNA from heads of first instar larvae were pooled and extracted using the Qiagen DNEasy Blood & Tissues kit (Qiagen). Total DNA quality was checked using the BioAnalyzer System (Agilent), and yield was quantified using a Qubit DNA HS Assay Kit (Thermo Fisher). DNA methylation libraries were constructed and sequenced (150bp paired-end) at a depth of 30x by Novogene (Hong Kong). The resulting sample group sizes for gene expression analysis were: 3-4 libraries for each group (pooled from the same families used for RNA-sequencing): FC_POP_FC_ENV_, FC_POP_NC_ENV_, NC_POP_NC_ENV_, NC_POP_FC_ENV_ (n=3-4/group; total of 14 libraries). Following initial QC (using FastQC v0.11.9), reads were trimmed using TrimGalore (0.6.4) to remove adaptor sequences and poor-quality reads. Bisulfite conversion efficiency across all samples was estimated by Novogene and was between 99.14-99.50% for all samples. Bisulfite-converted reads obtained from each library were mapped to the *N. Vespilloides* genome using Bismark (v0.22.3; Krueger & Andrews, 2011) and were quantified following deduplication of reads. Mapping rates for samples to the *N. vespilloides* reference genome were 65.45% ± 5.48, which after deduplication gave an average coverage of 17.78x ± 1.79 (mean ± SD).

### Differential Methylation Analysis

For differential methylation analysis of individual CpGs we first tested each site in every sample to determine sites that were significantly methylated using a binomial test. We then tested for differential methylation across sites using a weighted binomial glm (*p*-values were adjusted using Benjamini-Hochperg corrections for multiple testing). We then mapped CpGs to their feature using our custom genome annotation (see below for details) and bedtools (55). For analyses of gene body methylation, we collapsed across all CpG sites within each genic region (using bedtools) to compute the average weighted methylation value for each gene for each sample. Genes were clustered into methylated and unmethylated clusters using mixture models using the mixtools R package. To determine which genes were differentially methylated we used a weighted binomial GLM (base R stats package) and adjusted for multiple testing using the Benjamini-Hochberg procedure. Genes were considered differentially methylated if they had an adjusted *p*<0.01. Fishers’ exact tests for enrichment were run using the base R stats packages.

### Genome Annotation

To annotate exons in each genome we used existing annotations, excluding genes that were split across multiple contigs. To annotate regions which may contain promoters or enhancers (5’ UTR), we took 1,000 bases upstream of each gene, excluding genes where this exceeded the contig start or end point. We annotated introns based on the position of exons, excluding genes that were split across multiple contigs. To annotate TEs, we used RepeatModeller (v2.0.1) to generate a model of TEs for the *N. vespilloides* genome, and then RepeatMasker (v4.1.0) to annotate TEs based on the model for that genome.

## Data Access & Code Availability

All raw sequencing data generated have been submitted to the NCBI Gene Expression Omnibus (GEO) under accession number (GSE171776). All code for the analyses contained within this paper can be found at: https://github.com/r-mashoodh/nves_meth_evol.

## Acknowledgements

We would like to thank Benjamin Jarrett, Sue Aspinall, Darren Rebar, Ana Duarte and Matt Schrader for technical assistance with maintaining burying beetle populations. We would also like to thank Syuan-Jyan Sun, Samuel Lewis and the Miska lab for technical assistance with the molecular work. A special thanks to Benjamin Jarrett for constructive feedback and many helpful discussions. This project was supported by the European Research Council (310785 BALDWINIAN_BEETLES) and a Royal Society Wolfson Merit Award, both to R.M.K. The molecular work was supported by a Biotechnology and Biological Sciences Research Council Future Leaders Fellowship (BB/R01115X/1) to R.M.

## References

1. Holliday R. Epigenetics: a historical overview. Epigenetics. 2006 Jun;1(2):76–80.

2. Jaenisch R, Bird A. Epigenetic regulation of gene expression: how the genome integrates intrinsic and environmental signals. Nat Genet. 2003 Mar;33(S3):245–54.

3. Curley JP, Mashoodh R, Champagne FA. Epigenetics and the origins of paternal effects. Hormones and Behavior. 2011 Mar;59(3):306–14.

4. Miska EA, Ferguson-Smith AC. Transgenerational inheritance: Models and mechanisms of non-DNA sequence-based inheritance. Science. 2016 Oct 7;354(6308):59–63.

5. Sarkies P. Molecular mechanisms of epigenetic inheritance: Possible evolutionary implications. Seminars in Cell & Developmental Biology. 2020 Jan;97:106–15.

6. Mashoodh R, Habrylo IB, Gudsnuk KM, Pelle G, Champagne FA. Maternal modulation of paternal effects on offspring development. Proc R Soc B. 2018 Mar 14;285(1874):20180118.

7. Badyaev AV, Uller T. Parental effects in ecology and evolution: mechanisms, processes and implications. Phil Trans R Soc B. 2009 Apr 27;364(1520):1169–77.

8. Bonduriansky R, Day T. Extended heredity: a new understanding of inheritance and evolution. Princeton: Princeton University Press; 2018. 288 p.

9. Dixon G, Liao Y, Bay LK, Matz MV. Role of gene body methylation in acclimatization and adaptation in a basal metazoan. Proc Natl Acad Sci USA. 2018 Dec 26;115(52):13342–6.

10. Dubin MJ, Zhang P, Meng D, Remigereau M-S, Osborne EJ, Paolo Casale F, et al. DNA methylation in Arabidopsis has a genetic basis and shows evidence of local adaptation. Elife. 2015 May 5;4:e05255.

11. Heckwolf MJ, Meyer BS, Häsler R, Höppner MP, Eizaguirre C, Reusch TBH. Two different epigenetic information channels in wild three-spined sticklebacks are involved in salinity adaptation. Sci Adv. 2020 Mar;6(12):eaaz1138.

12. Liew YJ, Howells EJ, Wang X, Michell CT, Burt JA, Idaghdour Y, et al. Intergenerational epigenetic inheritance in reef-building corals. Nat Clim Chang. 2020 Mar;10(3):254–9.

13. Skinner MK, Gurerrero-Bosagna C, Haque MM, Nilsson EE, Koop JAH, Knutie SA, et al. Epigenetics and the Evolution of Darwin’s Finches. Genome Biology and Evolution. 2014 Aug;6(8):1972–89.

14. Hu J, Wuitchik SJS, Barry TN, Jamniczky HA, Rogers SM, Barrett RDH. Heritability of DNA methylation in threespine stickleback (Gasterosteus aculeatus). Peichel C, editor. Genetics. 2021 Mar 3;217(1):1–15.

15. Jarrett BJM, Evans E, Haynes HB, Leaf MR, Rebar D, Duarte A, et al. A sustained change in the supply of parental care causes adaptive evolution of offspring morphology. Nat Commun. 2018 Dec;9(1):3987.

16. Eggert A-K, Reinking M, Müller JK. Parental care improves offspring survival and growth in burying beetles. Animal Behaviour. 1998 Jan;55(1):97–107.

17. Kilner RM, Boncoraglio G, Henshaw JM, Jarrett BJM, De Gasperin O, Attisano A, et al. Parental effects alter the adaptive value of an adult behavioural trait. 2015;(4):e07340.

18. Schrader M, Jarrett BJM, Rebar D, Kilner RM. Adaptation to a novel family environment involves both apparent and cryptic phenotypic changes. Proc R Soc B. 2017 Sep 13;284(1862):20171295.

19. Schrader M, Jarrett BJM, Kilner RM. Using Experimental Evolution to Study Adaptations for Life within the Family. The American Naturalist. 2015 May;185(5):610–9.

20. Jarrett BJM, Rebar D, Haynes HB, Leaf MR, Halliwell C, Kemp R, et al. Adaptive evolution of synchronous egg-hatching in compensation for the loss of parental care. Proc R Soc B. 2018 Aug 29;285(1885):20181452.

21. Rebar D, Bailey NW, Jarrett BJM, Kilner RM. An evolutionary switch from sibling rivalry to sibling cooperation, caused by a sustained loss of parental care. Proc Natl Acad Sci USA. 2020 Feb 4;117(5):2544–50.

22. Law JA, Jacobsen SE. Establishing, maintaining and modifying DNA methylation patterns in plants and animals. Nat Rev Genet. 2010 Mar;11(3):204–20.

23. Arsenault SV, Hunt BG, Rehan SM. The effect of maternal care on gene expression and DNA methylation in a subsocial bee. Nat Commun. 2018 Dec;9(1):3468.

24. Franklin TB, Russig H, Weiss IC, Gräff J, Linder N, Michalon A, et al. Epigenetic Transmission of the Impact of Early Stress Across Generations. Biological Psychiatry. 2010 Sep;68(5):408–15.

25. Nyman C, Hebert FO, Bessert-Nettelbeck M, Aubin-Horth N, Taborsky B. Transcriptomic signatures of social experience during early development in a highly social cichlid fish. Mol Ecol. 2020 Feb;29(3):610–23.

26. Weaver ICG, Meaney MJ, Szyf M. Maternal care effects on the hippocampal transcriptome and anxiety-mediated behaviors in the offspring that are reversible in adulthood. Proceedings of the National Academy of Sciences. 2006 Feb 28;103(9):3480–5.

27. Bewick AJ, Vogel KJ, Moore AJ, Schmitz RJ. Evolution of DNA Methylation across Insects. Mol Biol Evol. 2016 Dec 25;msw264.

28. Cunningham CB, Ji L, Wiberg RAW, Shelton J, McKinney EC, Parker DJ, et al. The Genome and Methylome of a Beetle with Complex Social Behavior, Nicrophorus vespilloides (Coleoptera: Silphidae). Genome Biol Evol. 2015 Dec;7(12):3383–96.

29. Lewis SH, Ross L, Bain SA, Pahita E, Smith SA, Cordaux R, et al Widespread conservation and lineage-specific diversification of genome-wide DNA methylation patterns across arthropods. Reik W, editor. PLoS Genet. 2020 Jun 25;16(6):e1008864.

30. Feng S, Cokus SJ, Zhang X, Chen P-Y, Bostick M, Goll MG, et al. Conservation and divergence of methylation patterning in plants and animals. Proceedings of the National Academy of Sciences. 2010 May 11;107(19):8689–94.

31. Schmitz RJ, Lewis ZA, Goll MG. DNA Methylation: Shared and Divergent Features across Eukaryotes. Trends in Genetics. 2019 Nov;35(11):818–27.

32. Charlesworth D, Barton NH, Charlesworth B. The sources of adaptive variation. Proc R Soc B. 2017 May 31;284(1855):20162864.

33. Laland K, Uller T, Feldman M, Sterelny K, Müller GB, Moczek A, et al. Does evolutionary theory need a rethink? Nature. 2014 Oct;514(7521):161–4.

34. Gustafsson L, Sutherland WJ. The costs of reproduction in the collared flycatcher Ficedula albicollis. Nature. 1988 Oct;335(6193):813–5.

35. Steiger S. Bigger mothers are better mothers: disentangling size-related prenatal and postnatal maternal effects. Proc R Soc B. 2013 Sep 7;280(1766):20131225.

36. Massart R, Suderman M, Provencal N, Yi C, Bennett AJ, Suomi S, et al. Hydroxymethylation and DNA methylation profiles in the prefrontal cortex of the non-human primate rhesus macaque and the impact of maternal deprivation on hydroxymethylation. Neuroscience. 2014 May;268:139–48.

37. Wang L, Dankert H, Perona P, Anderson DJ. A common genetic target for environmental and heritable influences on aggressiveness in Drosophila. Proceedings of the National Academy of Sciences. 2008 Apr 15;105(15):5657–63.

38. Bentzur A, Shmueli A, Omesi L, Ryvkin J, Knapp J-M, Parnas M, et al. Odorant binding protein 69a connects social interaction to modulation of social responsiveness in Drosophila. Taghert PH, editor. PLoS Genet. 2018 Apr 9;14(4):e1007328.

39. Farhadian SF, Suárez-Fariñas M, Cho CE, Pellegrino M, Vosshall LB. Post-fasting olfactory, transcriptional, and feeding responses in Drosophila. Physiology & Behavior. 2012 Jan;105(2):544–53.

40. Suderman M, McGowan PO, Sasaki A, Huang TCT, Hallett MT, Meaney MJ, et al. Conserved epigenetic sensitivity to early life experience in the rat and human hippocampus. Proceedings of the National Academy of Sciences. 2012 Oct 16;109(Supplement_2):17266–72.

41. Yagound B, Remnant EJ, Buchmann G, Oldroyd BP. Intergenerational transfer of DNA methylation marks in the honey bee. Proc Natl Acad Sci USA. 2020 Dec 22;117(51):32519–27.

42. Feinberg AP, Irizarry RA. Stochastic epigenetic variation as a driving force of development, evolutionary adaptation, and disease. Proceedings of the National Academy of Sciences. 2010 Jan 26;107(suppl_1):1757–64.

43. Becker C, Hagmann J, Müller J, Koenig D, Stegle O, Borgwardt K, et al. Spontaneous epigenetic variation in the Arabidopsis thaliana methylome. Nature. 2011 Dec;480(7376):245–9.

44. Kawakatsu T, Huang SC, Jupe F, Sasaki E, Schmitz RJ, Urich MA, et al. Epigenomic Diversity in a Global Collection of Arabidopsis thaliana Accessions. Cell. 2016 Jul;166(2):492–505.

45. Cunningham CB, Ji L, McKinney EC, Benowitz KM, Schmitz RJ, Moore AJ. Changes of gene expression but not cytosine methylation are associated with male parental care reflecting behavioural state, social context and individual flexibility. J Exp Biol. 2019 Jan 15;222(2):jeb188649.

46. Pertea M, Kim D, Pertea GM, Leek JT, Salzberg SL. Transcript-level expression analysis of RNA-seq experiments with HISAT, StringTie and Ballgown. Nat Protoc. 2016 Sep;11(9):1650–67.

47. Love MI, Huber W, Anders S. Moderated estimation of fold change and dispersion for RNA-seq data with DESeq2. Genome Biol. 2014 Dec;15(12):550.

48. Jombart T. adegenet: a R package for the multivariate analysis of genetic markers. Bioinformatics. 2008 Jun 1;24(11):1403–5.

49. Hadfield JD. MCMC Methods for Multi-Response Generalized Linear Mixed Models: The MCMCglmm R Package. J Stat Soft [Internet]. 2010 [cited 2020 Jul 6];33(2). Available from: http://www.jstatsoft.org/v33/i02/

50. Stephens M. False discovery rates: a new deal. Biostat. 2016 Oct 17;kxw041.

51. Sun S, Catherall AM, Pascoal S, Jarrett BJM, Miller SE, Sheehan MJ, et al. Rapid local adaptation linked with phenotypic plasticity. Evolution Letters. 2020 Aug;4(4):345–59.

52. Huerta-Cepas J, Forslund K, Coelho LP, Szklarczyk D, Jensen LJ, von Mering C, et al. Fast Genome-Wide Functional Annotation through Orthology Assignment by eggNOG-Mapper. Molecular Biology and Evolution. 2017 Aug 1;34(8):2115–22.

53. Kriventseva EV, Kuznetsov D, Tegenfeldt F, Manni M, Dias R, Simão FA, et al. OrthoDB v10: sampling the diversity of animal, plant, fungal, protist, bacterial and viral genomes for evolutionary and functional annotations of orthologs. Nucleic Acids Research. 2019 Jan 8;47(D1):D807–11.

54. Krueger F, Andrews SR. Bismark: a flexible aligner and methylation caller for Bisulfite-Seq applications. Bioinformatics. 2011 Jun 1;27(11):1571–2.

55. Quinlan AR, Hall IM. BEDTools: a flexible suite of utilities for comparing genomic features. Bioinformatics. 2010 Mar 15;26(6):841–2.

